# SpheroScan: A User-Friendly Deep Learning Tool for Spheroid Image Analysis

**DOI:** 10.1101/2023.06.28.533479

**Authors:** Akshay Akshay, Mitali Katoch, Masoud Abedi, Mustafa Besic, Navid Shekarchizadeh, Fiona C. Burkhard, Alex Bigger-Allen, Rosalyn M. Adam, Katia Monastyrskaya, Ali Hashemi Gheinani

## Abstract

**Background:** In recent years, three-dimensional (3D) spheroid models have become increasingly popular in scientific research as they provide a more physiologically relevant microenvironment that mimics in vivo conditions. The use of 3D spheroid assays has proven to be advantageous as it offers a better understanding of the cellular behavior, drug efficacy, and toxicity as compared to traditional two-dimensional cell culture methods. However, the use of 3D spheroid assays is impeded by the absence of automated and user-friendly tools for spheroid image analysis, which adversely affects the reproducibility and throughput of these assays.

**Results:** To address these issues, we have developed a fully automated, web-based tool called SpheroScan, which uses the deep learning framework called Mask Regions with Convolutional Neural Networks (R-CNN) for image detection and segmentation. To develop a deep learning model that could be applied to spheroid images from a range of experimental conditions, we trained the model using spheroid images captured using IncuCyte Live-Cell Analysis System and a conventional microscope. Performance evaluation of the trained model using validation and test datasets shows promising results.

**Conclusion:** SpheroScan allows for easy analysis of large numbers of images and provides interactive visualization features for a more in-depth understanding of the data. Our tool represents a significant advancement in the analysis of spheroid images and will facilitate the widespread adoption of 3D spheroid models in scientific research. The source code and a detailed tutorial for SpheroScan are available at https://github.com/FunctionalUrology/SpheroScan.

**Key Points:**

- A deep learning model was trained to detect and segment spheroids in images from microscopes and Incucytes.
- The model performed well on both types of images with the total loss decreasing significantly during the training process.
- A web tool called SpheroScan was developed to facilitate the analysis of spheroid images, which includes prediction and visualization modules.
- SpheroScan is efficient and scalable, making it possible to handle large datasets with ease.
- SpheroScan is user-friendly and accessible to researchers, making it a valuable resource for the analysis of spheroid image data.

## Introduction

Two-dimensional (2D) cell culture models have long been a key component of biomedical research, but they often do not accurately replicate the in vivo environment [1]. In recent years, there has been an increasing realization that three-dimensional (3D) cell cultures, such as 3D spheroid models, are better able to mimic the in vivo environment. Moreover, the 3D cell cultures provide more clinically relevant insights into cellular behaviour and responses [2,3]. The 3D spheroid models, in particular, have become increasingly popular due to their ability to recreate the complex microenvironment found in vivo. This has made them a valuable tool for studying a variety of biological processes and diseases.

Tumour spheroids are widely used for testing anti-cancer medications [4]. They present a compromise between the cell accessibility of adherent cultures and the three-dimensionality of animal models. Spheroids retain more biological tumor features and reproduce intra-tumour environment, which is an important feature when selecting an effective treatment strategy. Most of the spheroid-based assays use the overall size and/or cell survival as a readout [5]. Thereby, a quick and easy tool for spheroid size estimation would be advantageous for such applications.

Another important area of research that is dependent on the spheroid size evaluation is the collagen gel contraction assay (CGCA) method [6]. CGCA is a widely used in vitro model for studying the interactions between cells and 3D extracellular matrices. These assays help understand matrix remodelling during fibrosis and wound healing. CGCA is a competent tool to evaluate the contractility of myofibroblasts harvested from fibrotic tissues. The advent of aqueous two-phase printing of cell-containing contractile collagen microgels has further advanced the CGCA technology [7]. Recently, the printing of the microscale cell-laden collagen gels has been combined with live cell imaging and automated image analysis to study the kinetics of cell-mediated contraction of the collagen matrix [8]. The image analysis method utilizes a plugin for FIJI, built around Waikato Environment for Knowledge Analysis (WEKA) Segmentation.

Despite the advantages of 3D spheroid models over 2D cell cultures, a major challenge has been the lack of fully automated and user-friendly tools for analysing spheroid images. This has hindered the widespread adoption of 3D spheroid models and has made high throughput analysis of spheroids difficult. Currently, the only options available for automatic spheroid detection in images are SMART [9] and SpheroidPicker [10]. However, SpheroidPicker is not open source, and using SMART requires a moderate to advanced level of computational and programming skills. As a result, many researchers with domain expertise are unable to utilize SMART easily. Furthermore, neither of these options provides visualization features to allow for efficient analysis of spheroid data. This is a significant drawback, as visualizing data can greatly aid in the interpretation and understanding of results.

To address these challenges, we have developed a fully automated, user-friendly web-based tool called SpheroScan for spheroid detection and interactive visualization of spheroid data using multiple publication-ready plots. Our tool is designed to be accessible to researchers regardless of their computational skills and aims to make the process of analysing spheroid images as simple and straightforward as possible. We have employed a state-of-the-art deep learning model called Mask R-CNN (Region-based Convolutional Neural Network) for image detection and segmentation. This model has proven to be highly effective in image analysis tasks and allows our tool to accurately detect and segment spheroids in images. With our tool, researchers can easily and quickly analyse large numbers of spheroid images and can use the interactive visualization features to gain a deeper understanding of their data (Figure 1).

**Figure 1.**
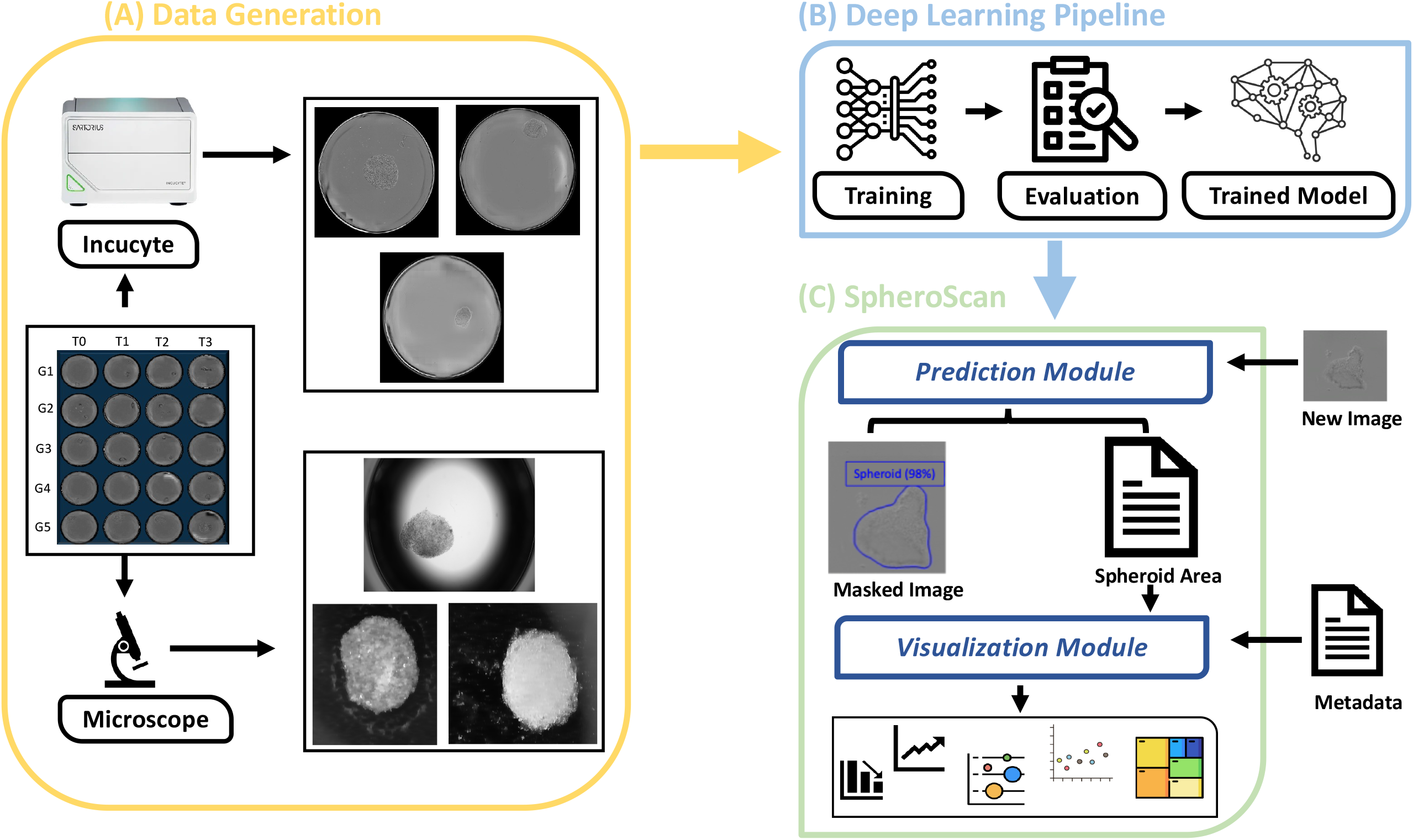
Graphical Abstract. **(A) Data Acquisition**. We used Incucyte and microscope platforms to generate spheroid images for the training and evaluation of deep learning model. **(B**) **Deep Learning (DL) Pipeline**. Two models were trained using Incucyte and microscope image datasets. These models were then evaluated on validation and test datasets. **(C) SpheroScan** consists of two submodules: Prediction and Visualization. The Prediction module applies the trained deep learning models to mask the input spheroid images, producing a CSV file with the area and intensity of each detected spheroid as output. The Visualization module enables the user to analyse the output from the Prediction module by providing various plots and statistical analyses.

## Results and Discussion

### Training and evaluating the performance of deep learning model

Figure 2 presents the performance of the trained Deep Learning (DL) model on the training, validation, and testing datasets for microscope and Incucyte images. The results show that the DL model was able to effectively learn and improve its performance over the course of training for both types of images. In particular, for Incucyte images, the total loss at baseline was 1.6 for the training data and 1.3 for the validation data. However, in the last epoch, the total loss reached its minimum values of 0.09 and 0.13 for the training and validation data, respectively (Figure 2A). This represents a significant improvement in performance. Similarly, the bounding box and mask loss started at relatively high values of 0.3 and 0.7, respectively, but decreased to their minimum values of 0.03 and 0.04 in the last epoch (Figure 2B). The model also performed well on the training and validation datasets for microscope images, with the total loss decreasing from 1.8 and 1.4 to 0.09 and 0.16 at the last epoch, respectively (Figure 2D). The bounding box and mask losses for the microscope dataset were also low, 0.036 and 0.045, respectively, at the last epoch (Figure 2E). Overall, these results demonstrate the robustness and effectiveness of the DL model in accurately detecting and segmenting spheroids in images from both microscopes and Incucytes.

**Figure 2.**
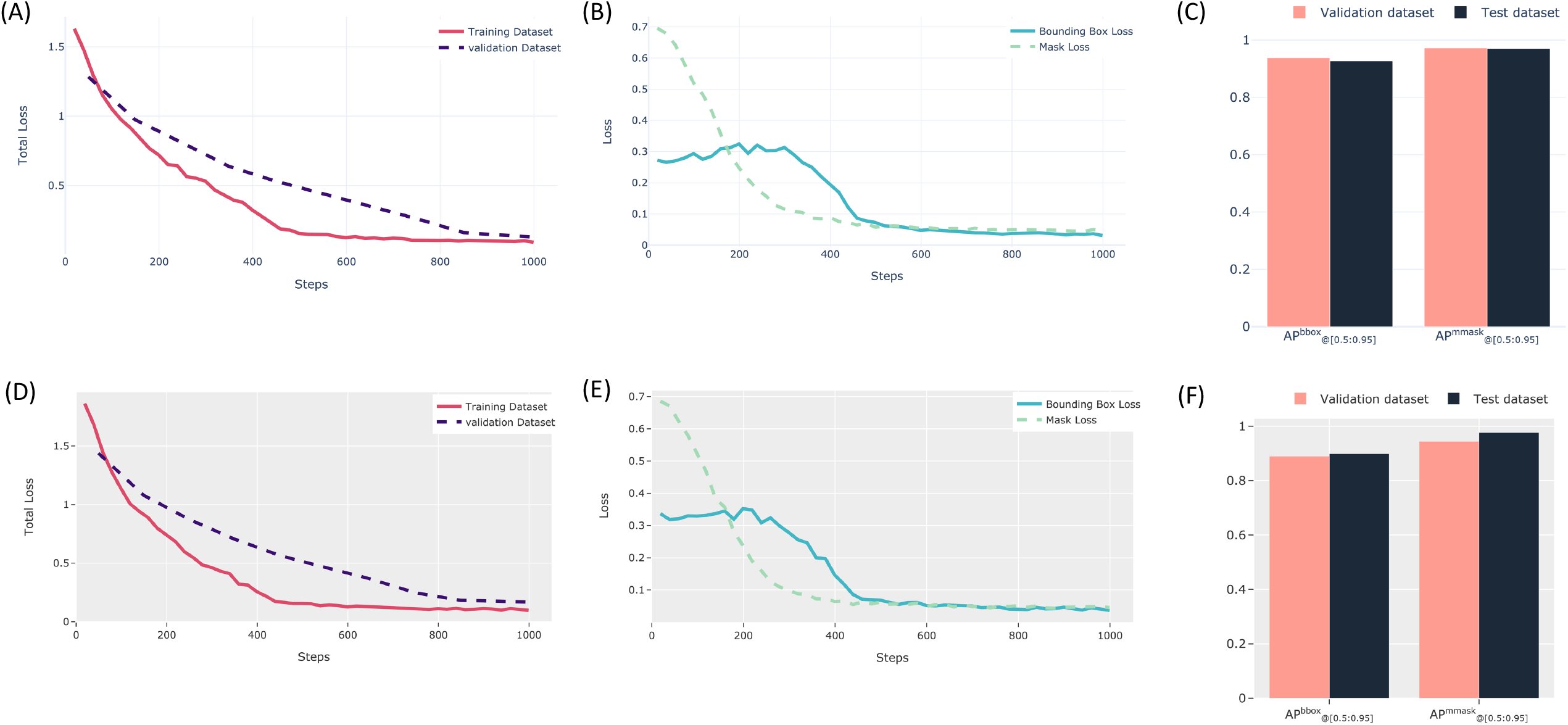
Results of the deep learning model’s performance. The total loss for both training and validation datasets of Incucyte **(A)** and microscope **(D)** images. The bounding box loss and mask loss for the training dataset of Incucyte **(B)** and microscope **(E)** images. The APbbox@_[0.5:0.95]_ and APmmask@_[0.5:0.95]_ for the validation and test datasets of Incucyte **(C)** and microscope **(F)** images. The APbbox@_[0.5:0.95]_ represents the average precision for bounding boxes, and the APmmask@_[0.5:0.95]_ represents the average precision for segmentation masks in the range of 0.5 to 0.95.

To evaluate the performance of the trained model in segmenting spheroids, we calculated the Average Precision (AP) metric for bounding boxes and segmentation masks in the range of 0.5 to 0.95. Throughout the text, APbbox@_[0.5:0.95]_ represents the AP for bounding boxes, and APmmask@_[0.5:0.95]_ represents the AP for segmentation masks. In general, the trained models showed similar performance on the test and validation datasets. The values for APbbox@_[0.5:0.95]_ and APmmask@_[0.5:0.95]_ were 0.937 and 0.972, respectively, for the validation data, and 0.927 and 0.97, respectively, for the test data of Incucyte images (Figure 2C). The model’s performance on the validation and test datasets for microscopic images were also strong, with scores of 0.89 and 0.944 for APbbox@_[0.5:0.95]_ and APmmask@_[0.5:0.95]_ respectively on the validation data, and scores of 0.899 and 0.977 respectively on the test data (Figure 2F).

### SpheroScan characteristics

We have developed an open-source web tool called SpheroScan to facilitate the analysis of spheroid images. This user-friendly, interactive tool is designed to streamline the process of spheroid segmentation, area calculation, and downstream analysis of spheroid image data. Furthermore, it helps to standardize and accelerate the analysis of spheroid assay results. SpheroScan consists of two main modules: prediction and visualization. The prediction module uses previously trained DL models to detect the spheroid in the input images; accordingly, a CSV file is generated with the area and intensity of each detected spheroid (Figure S1.A). The visualization module allows the user to analyse the results of the prediction module through various types of plots and statistical analyses (Figure S1.B). The plots generated by the visualization module are ready for publication and can be saved as high-quality images in PNG format. Overall, SpheroScan is a powerful and user-friendly tool that greatly simplifies and enhances the analysis of spheroid image data (Figure S2-S4).

The runtime complexity of the prediction module is linear, meaning that it scales in proportion to the size of the input data. This is an important property because it means that the prediction module will be efficient and scalable, even when processing large datasets. To confirm the linear runtime complexity of the prediction module, we tested it on four different image datasets with various numbers of images. The results of these tests showed that the prediction module consistently had a linear runtime, taking less than one second to mask a single image (Figure 3D). This demonstrates that the prediction module is highly efficient and capable of handling large datasets with ease. We evaluated the run-time performance on a Red Hat server with 16 Central Processing Unit (CPU) cores and 64 GB of Random-Access Memory (RAM).

**Figure 3.**
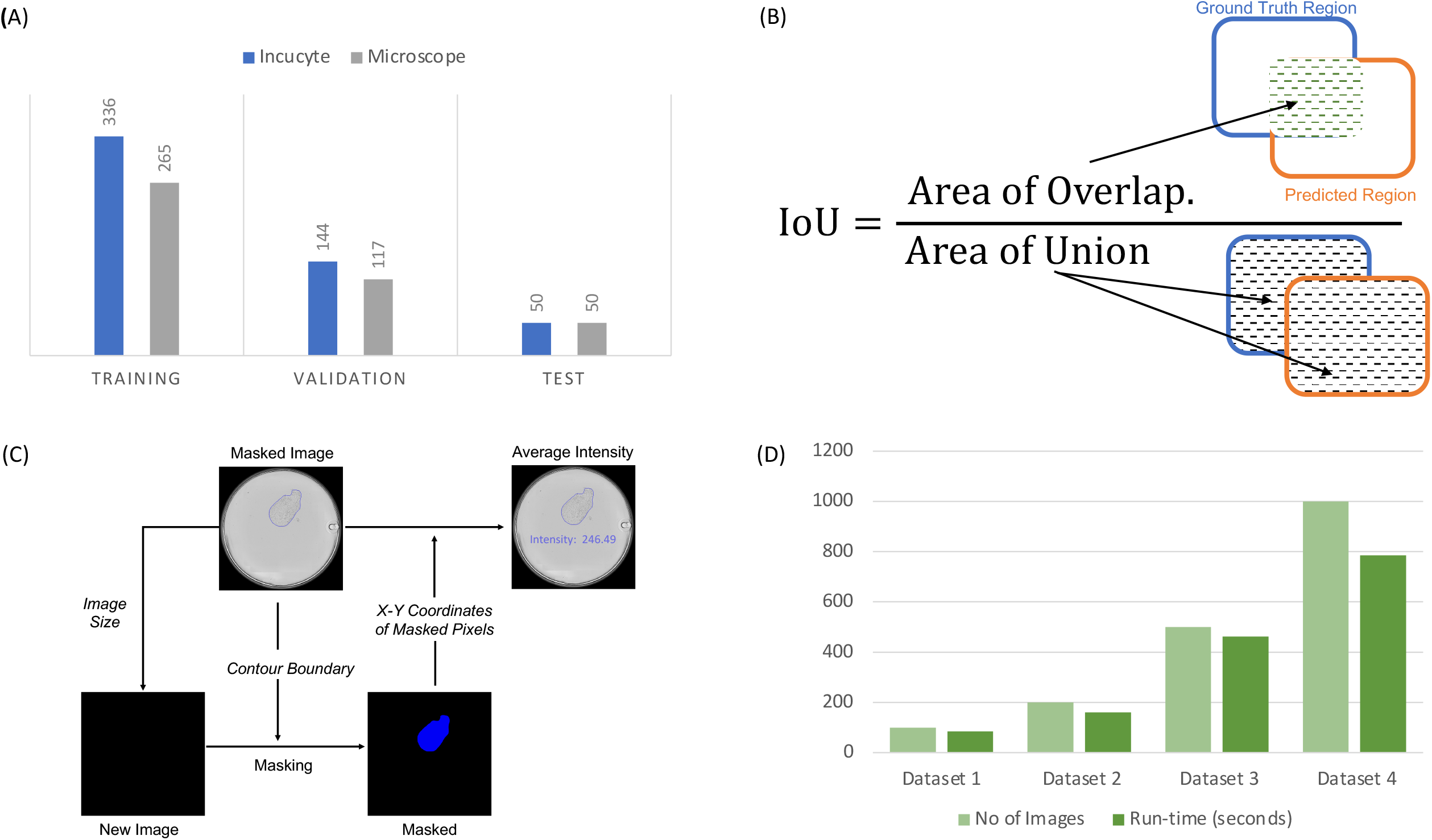
**(A) Datasets.** Training, validation and test datasets size for Incucyte and microscpe model. **(B) IoU metric**. The Intersection over Union (IoU) metric is a measure of the overlap between two bounding boxes or masks. It is calculated by dividing the overlap area between the predicted and ground truth regions by the total area of both regions combined. **(C) Spheroid Intensity calculation**. To determine the intensity of the spheroid image, a new image with the same shape and number of pixels as the original is created, but with all pixels set to zero intensity. The predicted contour boundary of the spheroid or spheroids is applied to this new image, and all pixels inside the boundary are set to 255 intensity. The x and y coordinates of each pixel in the new image with a value of 255 are then extracted. The average pixel intensity value for all points within the contour boundary is then calculated using Python’s OpenCV module. **(D) Run-time analysis**. The runtime complexity of the prediction module was analysed using four different image datasets of varying sizes. The results showed that it takes less than a second to mask an image, and the runtime complexity of the prediction module is linear. This means that the time required to process an image increases in proportion to the number of images being processed.

### Limitations and considerations

As with any technology, there are limitations and considerations to keep in mind when using the SpheroScan system. First, it is important to note that this developed tool is primarily designed for use with the spheroid images from Incucyte and microscope platforms. Additionally, when analyzing images that contain more than one spheroid, the performance of the SpheroScan system may decrease. Therefore, it is important to carefully consider the experimental design and imaging conditions to ensure optimal performance and accurate results. The authors aim to expand the training dataset with a diverse range of external images from various experimental environments and platforms in the future to improve and advance the utility of SpheroScan. Generally, while the SpheroScan system offers many advantages for high-throughput spheroid analysis, it is important to be aware of its limitations and take steps to address them as needed.

### Conclusion

The development of the web-based tool SpheroScan represents a significant advancement in the analysis of 3D spheroid images. Using the state-of-the-art deep learning techniques, our tool accurately detects and segments spheroids in images, making it easy for researchers to analyse large numbers of spheroid images. Additionally, our tool is user-friendly and accessible to researchers regardless of their computational skills, making it a valuable resource for the scientific community. The interactive visualization features provided by our tool also allow for a more in-depth understanding of spheroid data, which will further facilitate the widespread adoption of 3D spheroid models in research. Overall, SpheroScan represents a valuable tool for researchers working with 3D spheroid models and will help to advance the use of these models in scientific research.

## Materials and Methods

### Availability and implementation

The source code, example input data, and a detailed tutorial for SpheroScan are available at https://github.com/FunctionalUrology/SpheroScan. SpheroScan was developed using Plotly Dash [11] library in Python (version 3.10.6) and all the plots were made using Plotly. Pandas library [12,13] was used to store and process the data.

### Spheroid image acquisition

In this study, our goal was to create a generalized DL model that can be used for spheroid images from various experimental setups or laboratory environments. To this end, we applied the aqueous two-phase solution method [7] to embed the cells of interest into collagen matrix spheroids. To estimate the cell-driven contraction of the collagen matrix, we collected spheroid images from different treatment conditions and time points, using both bladder Smooth Muscle Cells (SMCs) and Human Embryonic Kidney (HEK) cells. SMC cells were chosen for this study since they have the ability to contract, which we expected to lead to the creation of spheroids in a wide range of sizes. HEK cells, on the other hand, do not contract and were used as a negative control to ensure the accuracy of our results. The spheroids were treated with various concentrations of histamine and Fetal Bovine Serum (FBS) and were observed at regular intervals to track their response to these treatments.

To generate the image datasets needed for a DL model, we performed a spheroid gel contraction assay using 5000 SMC or HEK cells per collagen spheroid. After the collagen droplet polymerized, the medium was changed and plates were transferred to an Incucyte Live-Cell Analysis System, which acquired images of the spheroids every hour for 24 hours. Alternatively, we used a ZEISS Axio Vert.A1 Inverted Microscope and manually acquired images of the spheroids at selected time points. By using both methods, we were able to capture a wide range of spheroid images and to create a robust dataset for our DL model.

A total of 480 images were obtained from the Incucyte system, and these were randomly divided into a training dataset of 336 images (70%) and a validation dataset of 144 images (30%). An additional test dataset of 50 images was used to evaluate the performance of the trained model. To create a model specifically for microscopic images, we gathered spheroid images from the microscope and divided them into three datasets: training, validation, and test. The training dataset included 265 images, the validation dataset included 117 images, and the test dataset included 50 images (Figure 3A). To test the robustness of the trained model, the spheroids in the test dataset were treated differently from those in the training and validation datasets. The medium used here was smooth muscle cell medium and Dulbecco’s Modified Eagle Medium (DMEM) with 0.5% and 1% FBS.

In the following step, an experienced researcher in the spheroid assay manually annotated the images from Incucyte and microscopes using the VGG Image Annotator [14].

### Deep learning framework

For spheroid detection and segmentation, we used a state-of-the-art DL model called Mask R-CNN and an open-source Python [15] library called Detectron2 [16]. Mask R-CNN is a method for solving the problem of instance segmentation, which involves both object detection and semantic segmentation. Object detection is the process of identifying and classifying multiple objects within an image, while semantic segmentation involves understanding the image at the pixel level to distinguish individual objects within the image. In order to perform these tasks, Mask R-CNN first uses a deep Convolutional Neural Network (CNN) to process the input image and to generate a set of feature maps. These feature maps are then used as input for the next step in the process.

Mask R-CNN performs object detection in two stages. First, it uses a Region Proposal Network (RPN) module to identify Regions of Interest (ROIs) within the image. ROIs are defined as bounding boxes with a high probability of containing objects. In the second stage, Mask R-CNN uses an ROI classifier and bounding box regressor module to classify the objects within the ROIs and to determine their bounding boxes. Both the RPN and ROI classifier and bounding box regressor modules are implemented as CNNs.

For semantic segmentation, Mask R-CNN uses a fully convolutional network (FCN) called the mask segmentation module to predict masks for each ROI determined in the object detection phase. This allows Mask R-CNN to accurately identify and distinguish individual objects within the image and segment them from the background. Overall, the combination of object detection and semantic segmentation allows Mask R-CNN to achieve highly accurate and detailed instance segmentation results (Figure S5).

In this study, we used the Mask R-CNN model for instance segmentation and tuned several of its parameters to fit the specific problem and the dataset we were working with. The backbone of the model was a ResNet-50 feature pyramid network, and we initialised the model with weights from a pre-trained COCO instance segmentation model. The batch size for training was set to 4, and the base learning rate was set to 0.00025. The RoIHead batch size was 256, and we used a single output class (for spheroids). We trained the model for a total of 1000 iterations. In addition to these specified parameters, we used the default values for all other parameters of the Mask R-CNN model.

### Evaluation Metrics

To evaluate the performance of the trained models on spheroid segmentation, we used the Average Precision (AP) or Mean Average Precision (mAP) metric. mAP is a commonly used evaluation metric in computer vision for measuring the accuracy of instance segmentation and object detection models. Many of the state-of-the-art object detection algorithms, such as Faster R-CNN [17], Mask R-CNN [18], MobileNet SSD [19], and YOLO [20], and benchmark challenges such as PASCAL VOC [21], use AP to evaluate their models. Calculation of AP is dependent on the following metrics:

#### Precision

It is defined as the fraction of true instances among all predicted instances and is calculated using the following formula:

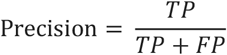

#### Recall

It is a metric that represents the fraction of retrieved instances among all relevant instances and is calculated as follows:

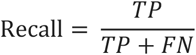

#### Intersection over Union (IoU)

The intersection over union (IoU) is a metric that measures the overlap between two bounding boxes or masks. It is commonly used to evaluate the accuracy of object detection and instance segmentation models. The IoU value ranges from 0 to 1, with a value of 1 indicating a completely accurate prediction. To calculate the IoU, the overlap between the predicted and ground truth regions is first determined and divided by the total area of both regions. The IoU is a useful metric because it allows for comparing predictions with different shapes and sizes, as it considers the area of both the predicted and ground truth regions (Figure 3B).

#### Average Precision (AP)

The Average Precision (AP) is a metric used to evaluate the performance of object detection and instance segmentation models. It is calculated as the area under the precision-recall curve, which plots the precision (the proportion of true positive detections among all positive detections) against the recall (the proportion of true positive detections among all ground truth objects) of a model. AP ranges from 0 to 1, with a higher value indicating better performance. A higher AP value indicates that the model can achieve both high precision and high recall, making it a useful metric for evaluating the overall performance of a model. AP can be calculated for a specific IoU threshold as follows:

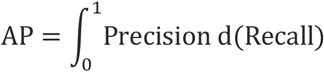

Often, AP is used as the average over multiple IoU thresholds, and it is calculated as follows:

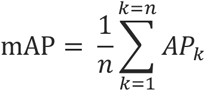

where,

*Ap*_*K*_= AP at *k*^th^ IoU threshold

*n* = Number of IoU thresholds under consideration.

In the following, AP_@0.75_ represents AP at IoU threshold 0.75 and AP_@[0.5:0.95]_ represents the average AP over 10 IoU thresholds (from 0.5 to 0.95 with a step size of 0.05).

### Area and Intensity Calculation

After performing object detection and instance segmentation on an image, we can use the predicted contour boundary of each spheroid to calculate its area and intensity. To calculate the area of a spheroid, we use Python’s OpenCV library to count the number of pixels within the contour boundary. This gives us the total area of the spheroid in pixels. To calculate the intensity of the spheroid, we follow a similar process. First, we create a new image with the same shape and number of pixels as the original, but with a default intensity of zero. This image is then masked with the predicted contour boundary of the spheroid, setting all pixels within the boundary to a value of 255. We then extract the *x* and *y* coordinates of all pixels with a value of 255, which correspond to the pixels within the contour boundary of the spheroid in the original image. Finally, we use OpenCV to calculate the average intensity of these pixels, which gives us the intensity value for the spheroid. This process allows us to accurately measure the area and intensity of each spheroid in an image (Figure 3C).

## Availability of supporting source code and requirements

Project name: SpheroScan

Project home page: https://github.com/FunctionalUrology/SpheroScan

Operating system(s): Linux or Mac

Programming language: Python 3.10.6

Other requirements: Docker, Python, Anaconda, Git License: GNU GPL

## Author contributions statement

K.M., A.H.G, and A.A. conceived the idea for the manuscript. M.B generated all the data. A.A. developed the deep learning pipeline. A.A and M.K developed the code for SpheroScan. K.M., F.C.B, and A.H.G tested the SpheroScan and provided scientific inputs throughout the development phase. F.C.B, R.M.A and A.B.A provided the feedback on biological application of the tool. N.S and M.A provided the mathematical support and did the testing and debugging. All authors contributed to writing, proofreading, and correcting the manuscript.

## Data Availability

All supporting data, which includes images used for training, validation, and testing [22], as well as the trained model weights [23], is available at zenodo.

## Acknowledgments

We are thankful to Ankush Sharma for his guidance and to Niharika Jakhar for testing SpheroScan on various operating systems.

