## Supplementary file for "SpheroScan: A User-Friendly Deep Learning Tool for Spheroid Image Analysis"

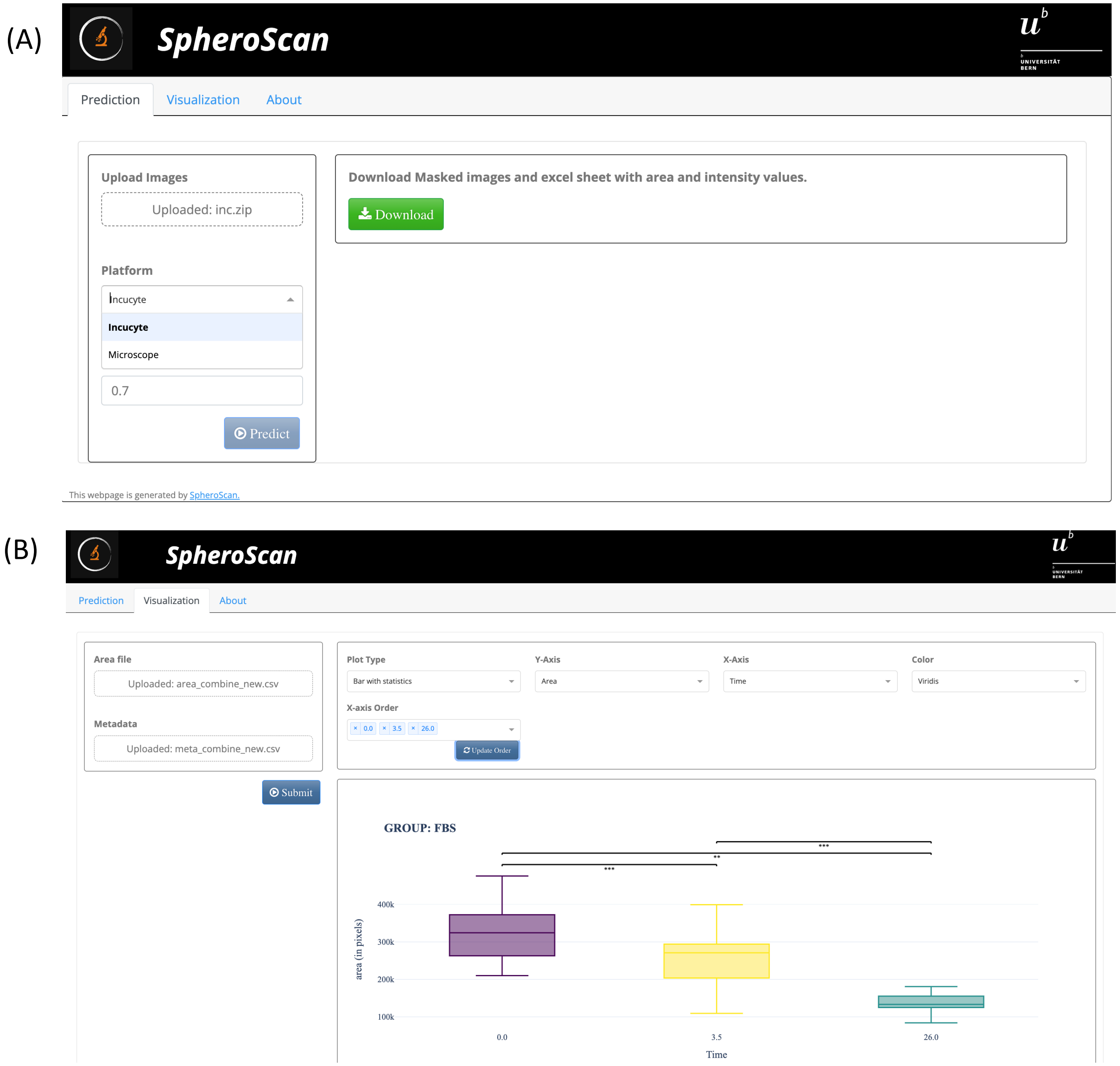


**Figure S1. SpheroScan Graphical User Interface. (A) Prediction Module.** The Prediction Module applies trained DL models to identify and mask spheroid images. It requires a zipped folder of images, platform type, and prediction threshold as input and generates masked images and a CSV file containing the area and intensity data of the identified spheroids as output. **(B) Visualization Module.** The Visualization Module creates plots and performs statistical analysis using the output file from the Prediction Module and a metadata file that contains information about the study design. It offers various types of plots and allows users to customize the plot options, such as plot type and color palette. Users can export plots in high-resolution PNG format.

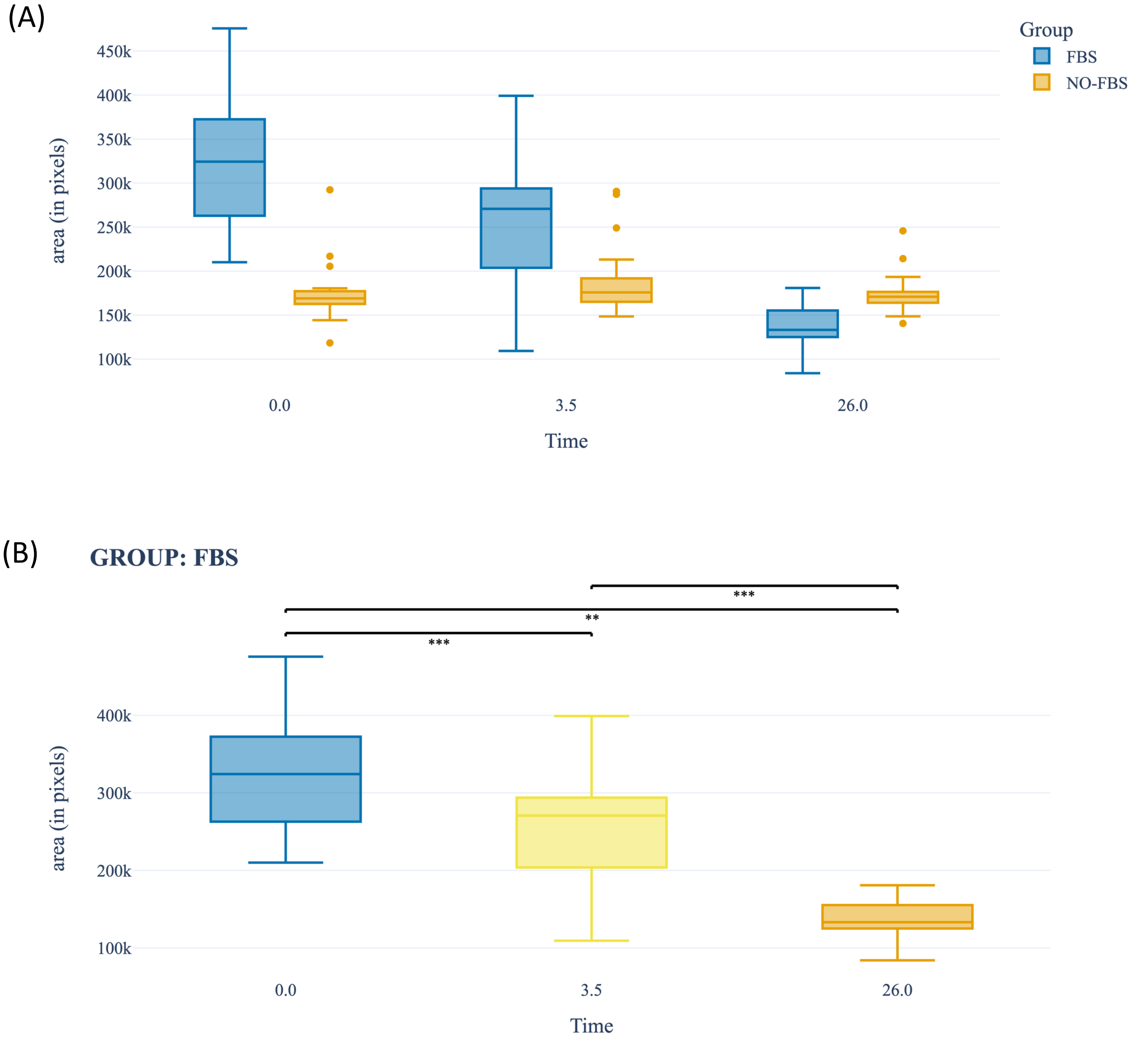


**Figure S2. SpheroScan Plot Gallery. (A)** Bar Plot **(B)** Bar Plot with significance level. A bar plot with significance level is a visual representation of data where the level of significance is indicated by stars. Three stars (***) indicate a p-value of less than 0.001, while "ns" represents a p-value of 0.05 or greater. The less stars, the lower the significance level.

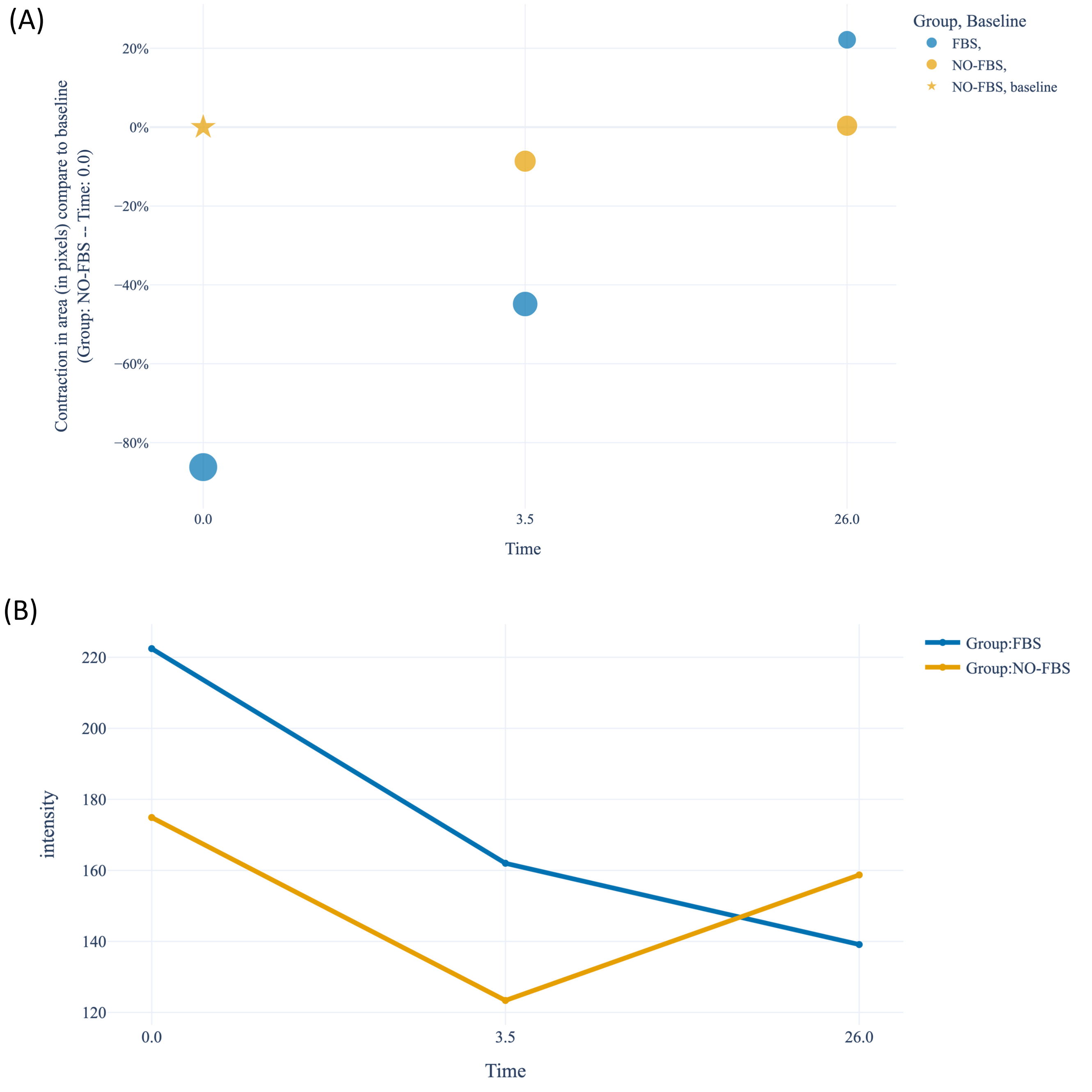


**Figure S3. SpheroScan Plot Gallery. (A)** Bubble Plot. A bubble plot is a type of scatter plot where the size of the bubbles represents the mean spheroid area for a certain group. The Y-axis displays the relative area or contraction of the spheroid, calculated with respect to a baseline group. **(B)** Line plot

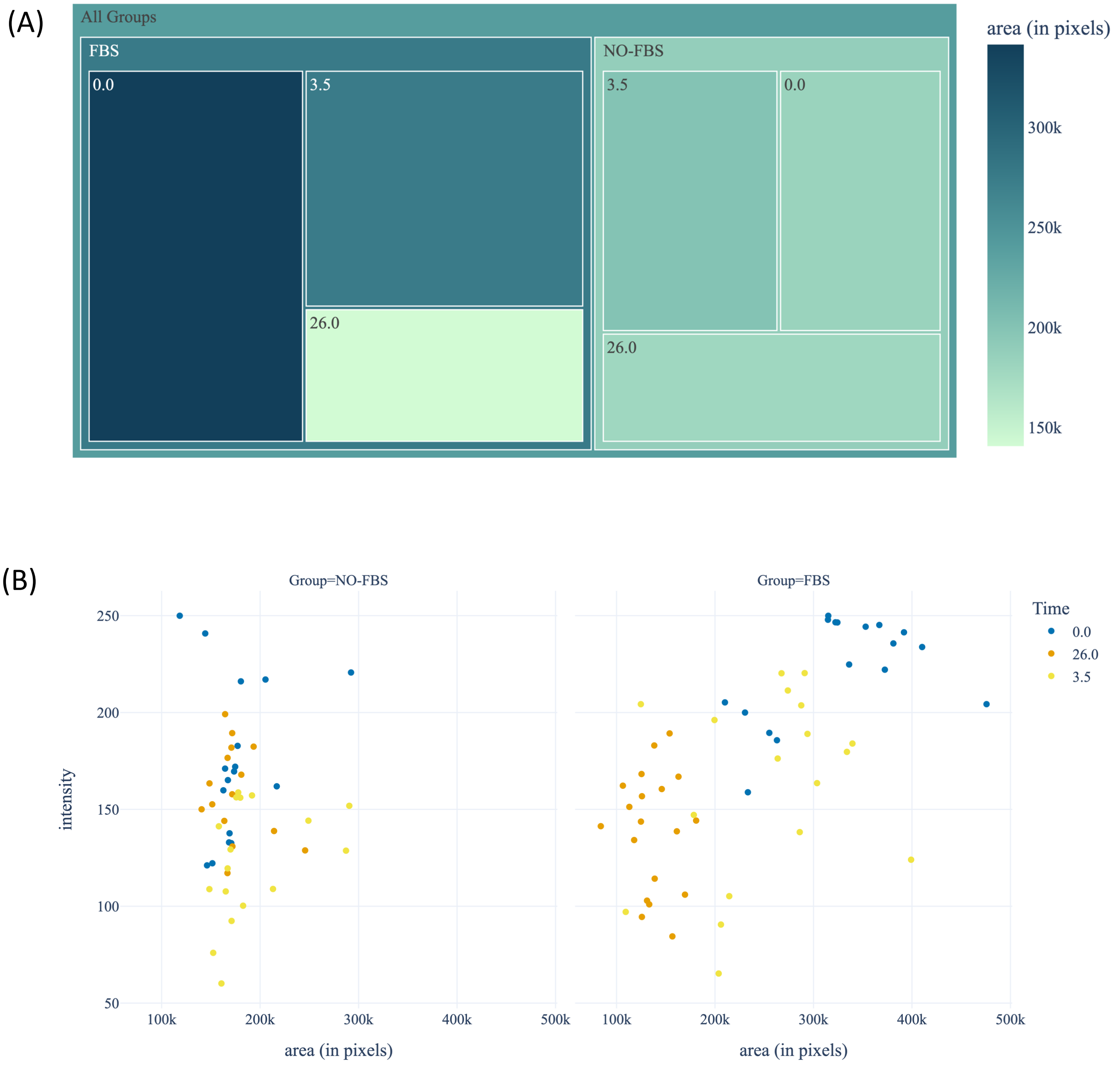


**Figure S4. SpheroScan Plot Gallery. (A)** Treemap. A treemap is a method of displaying hierarchical data in which nested rectangles are used to represent different groups. The outer rectangles represent the top-level groups, while the inner rectangles represent sub-groups. The size and color of each rectangle in the treemap indicate the mean spheroid areas or intensity of the corresponding group. **(B)** Scatter plot

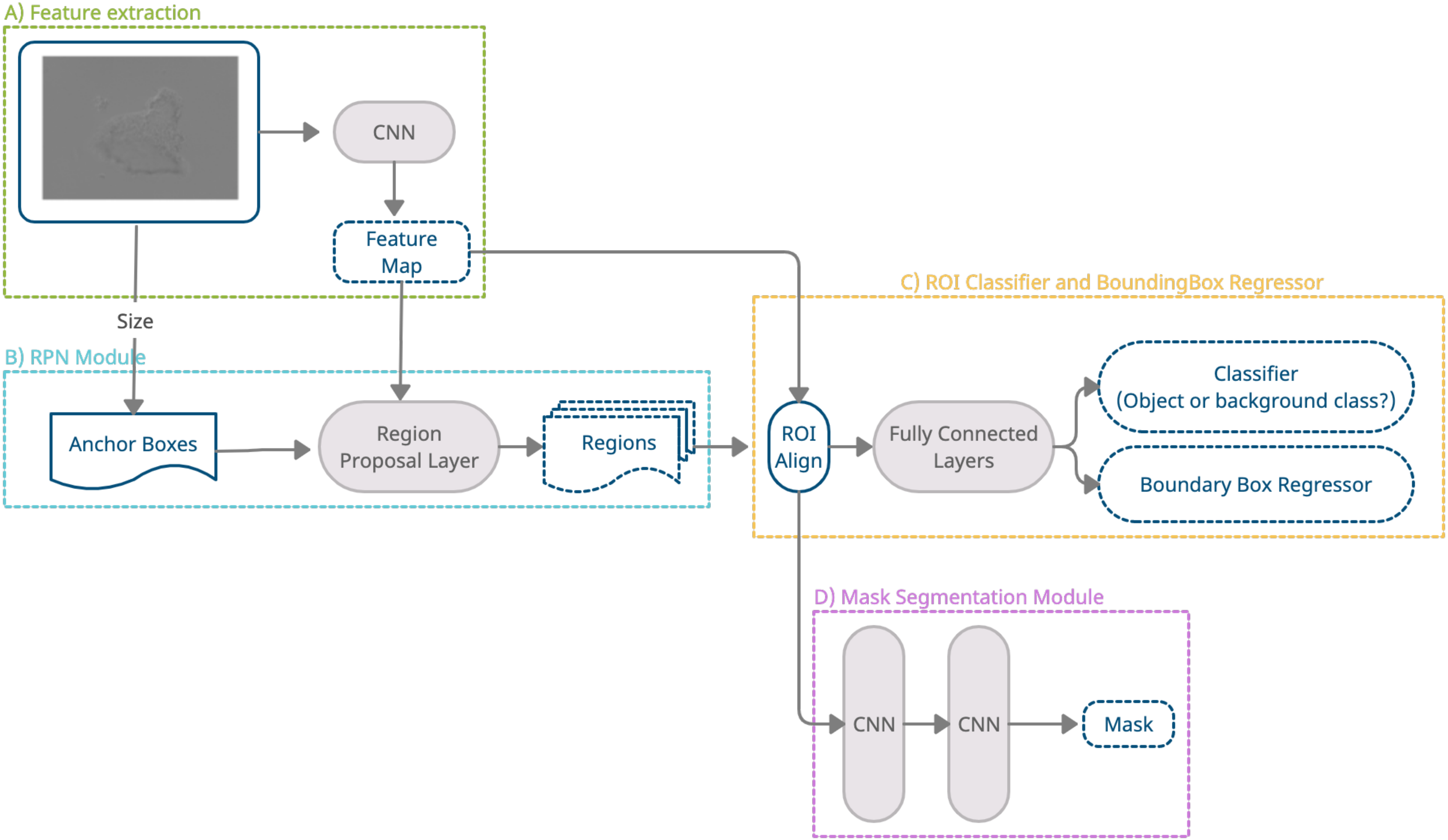


**Figure S5. Mask R-CNN Architecture.** The Mask R-CNN model consists of four main modules: feature extraction, Region Proposal Network (RPN), Region of Interest (ROI) classifier and bounding box regressor, and mask segmentation. The feature extraction module takes images as input and produces feature maps. The RPN module then runs on the feature maps and uses a sliding window to identify bounding boxes with a high likelihood of containing objects (ROIs). For each ROI, the ROI classifier and bounding box regressor module is used to determine the class label of the object. For semantic segmentation, the Mask R-CNN model uses a Fully Convolutional Network (FCN) in the mask segmentation module to predict a mask for each ROI identified in the object detection phase.
